# Visualizing the Role of Lipid Dynamics during Infrared Neural Stimulation with Hyperspectral Stimulated Raman Scattering Microscopy

**DOI:** 10.1101/2021.05.24.444984

**Authors:** Wilson R Adams, Rekha Gautam, Andrea Locke, Ana I. Borrachero-Conejo, Bryan Dollinger, Graham A. Throckmorton, Craig Duvall, E Duco Jansen, Anita Mahadevan-Jansen

**Affiliations:** Dept. of Biomedical Engineering, Vanderbilt University, Nashville, TN, USA; Dept. of Neurosurgery, Vanderbilt University Medical Center, Nashville, TN, USA

## Abstract

Infrared neural stimulation, or INS, is a method of using pulsed infrared light to yield label-free neural stimulation with broad experimental and translational utility. Despite its robust demonstration, the mechanistic and biophysical underpinnings of INS have been the subject of debate for more than a decade. The role of lipid membrane thermodynamics appears to play an important role in how fast IR-mediated heating nonspecifically drives action potential generation. Direct observation of lipid membrane dynamics during INS remains to be shown in a live neural model system. To directly test the involvement of lipid dynamics in INS, we used hyperspectral stimulated Raman scattering (hsSRS) microscopy to study biochemical signatures of high-speed vibrational dynamics underlying INS in a live neural cell culture model. Findings suggest that lipid bilayer structural changes are occurring during INS *in vitro* in NG108-15 neuroglioma cells. Lipid-specific signatures of cell SRS spectra were found to vary with stimulation energy and radiant exposure. Spectroscopic observations were verified against high-speed ratiometric fluorescence imaging of a conventional lipophilic membrane structure reporter, di-4-ANNEPS. Overall, the presented data supports the hypothesis that INS causes changes in the lipid membrane of neural cells by changing lipid membrane packing order – which coincides with likelihood of cell stimulation. Furthermore, this work highlights the potential of hsSRS as a method to study biophysical and biochemical dynamics safely in live cells.

## Main Text

Neuromodulation using directed energy, including optical, ultrasonic, and radio frequency, have gained notable interest recently due to their spatial precision, noninvasive implementation, and promising potential for clinical translation in therapeutic interventions. Label-free optical neuromodulation with pulsed infrared (IR) light, or infrared neural stimulation (INS), offers a spatially and temporally precise means of contact-free activation of neural cells without the need for genetic modification or exogenous mediators. Similar to most label-free directed energy methods of neuromodulation, the biophysical mechanisms underlying INS have remained elusive for more than a decade (1). In contrast to the tools derived from molecular biology, such as optogenetics or photochemical uncaging, INS appears to act through an entirely different photothermal-based mechanism (1, 2). The role of lipid membrane dynamics are thought to play an important role in how IR light depolarizes neurons photothermally (3), but remains to be directly experimentally validated in a live neural model system.

Infrared wavelengths generally used for INS are strongly absorbed by water (4, 5). The rapid temperature rise from brief pulses of IR light were experimentally shown to depolarize HEK cells as well as synthetic charged lipid bilayer preparations through a transient increase in membrane capacitance (2). Biomolecular explanations for these observations were unclear. A biophysical explanation of this phenomenon was described computationally by factoring in the thermal dependence of lipid bilayer geometry with a Gouy-Chapman-Stern based electrodynamic model of charged lipid bilayers (3). While the experimental data and computational model agree with each other, the role of lipid dynamics in neural models of INS remain to be directly validated. Lipid dynamics during INS have been probed through electrophysiology and fluorescent membrane structure reporters (2, 6, 7). However, these methods are inherently indirect to lipid molecular dynamics. There remains to be any direct observation of lipid dynamics in live neural cells during INS. Understanding the role of lipid dynamics in the mechanisms of INS would provide both valuable scientific insight and a basis for innovation towards the next generations of neuromodulation technology.

Conventional methods of directly measuring lipid bilayer geometry, such as x-ray diffraction and small angle neutron scattering, are slow and not biologically compatible (8–10). Optical methods are well suited for high resolution, biologically compatible experiments, but generally lack the spatial resolution necessary to directly resolve lipid bilayer geometry (< 3 nm thick) on millisecond timescales. Fluorescent functional lipid indicators, such as laurdan or di-4-ANNEPS (11), have been shown to be powerful tools in studying lipid membrane biophysics. However, these indicators offer latent readouts of lipid dynamics and are inherently indirect in that they rely on the molecular interaction of reporter molecules with their molecular environment rather than the lipid molecules themselves. Vibrational spectroscopic methods, such as Raman scattering and infrared absorption, can be performed label-free and offer a feature-rich molecular signature useful in studying lipid organization in live cells. Traditionally, vibrational spectroscopic methods have not been biologically compatible on sub-second timescales (12, 13).

Stimulated Raman scattering (SRS) microscopy combines label-free vibrational spectroscopic contrast with subcellular spatial resolution and sub-second temporal resolution enabling time resolved vibrational spectral measurements of live neural cells during INS (14, 15). Others have shown that lipid molecular symmetry and ordered molecular interactions of water with lipid bilayers are observable with nonlinear Raman microscopy (16, 17). Moreover, SRS can be implemented fast enough to discern signatures of neuronal action potentials at millisecond timescales (16–18). With this in mind, we set out to employ a hyperspectral SRS (hsSRS) microscopy approach to identify vibrational signatures of lipid bilayer dynamics during INS in live neural cell cultures.

The goal of this paper is therefore to identify the molecular dynamics of membrane lipids in live neural cells during INS with hsSRS microscopy. We demonstrate a time-resolved hsSRS methodology combined with focus precompensation to obtain SRS spectra of live NG108 cells. Spectra of NG108 cells show significant changes during INS which are attributable to changes in lipid packing order and solvent interactions. Validation of this approach was compared to gold-standard ratiometric fluorescence of a functional lipid packing order indicator – di-4-ANNEPS. We discuss how changes in the vibrational spectral signatures of cells during INS compare to what the current mechanistic hypothesis implies. Furthermore, we offer practical insight to performing high-resolution optical microscopy in dynamically varying optical imaging conditions during INS, as well as offer some thoughts to the potential extensions of this hsSRS methodology as SRS technology continues to develop.

## Methods

### Cell Culture and Maintenance

Methods for neuronal hybridoma cell cultures were adapted from previous work (19, 20). A spiking neuroma-glioblastoma hybridoma cell line, NG-108-15 (Sigma-Aldrich, St. Louis, MO), were thawed and maintained in culture for 1 week prior to imaging experiments. Cells were maintained in Dulbecco’s Modified Eagle Medium supplemented with 4.5g/L of glucose, 20mM of L-glutamine, 15%v/v fetal bovine serum and 1%v/v of penicillin/streptomycin antibiotics. Cells were incubated at 37°C in 5% of CO_2_ and 95% relative humidity. Growth medium was completely replaced every 48 hours until cells approached confluency. Once ~80% confluent, cells were mechanically dissociated and propagated onto additional cell culture flasks until experimental use. All cells were imaged within 15 rounds of passage from thawed supplier stocks. Seventy-two hours prior to imaging, cells were passaged and plated onto poly-D-lysine-coated glass-bottom petri dished (Mattek, Natick, MA) to allow for cellular adherence. Twenty-four hours prior to image experiments, the cell culture medium was replaced with an identical DMEM formulation except for the reduction of fetal bovine serum concentration (3%v/v) to promote morphological differentiation into dendritic neuronal phenotypes. During imaging experiments, cells were maintained at room temperature and humidity in neurophysiologically balanced saline free of protein and glucose with the following composition (in mM): 140 NaCl, 4 KCl, 2 MgCl_2_, 2 CaCl_2_, 10 HEPES, 5 glucose, pH 7.4 with NaOH and osmolarity adjusted to ~318 mOsm with mannitol (21). Cells were imaged for 45 minutes before being discarded.

### Microscope System

The physical layout and capability of the custom-built multimodal imaging platform utilized in this study (**Figure 1A**) was described previously (22). Briefly, a dual output femtosecond near-infrared laser source (Insight DS+, Spectra Physics, Santa Clara, CA) was used to excite nonlinear contrast. Both output beams were spatially and temporally combined, with 20 MHz intensity modulation of the 1040 nm output and a variable linear optical path length on the 798 nm output for temporal collinearity and to facilitate hyperspectral SRS (23). The combined ultrafast laser outputs were subsequently chirped through 150mm of high-index SF11 glass rods (Newlight Photonics, Ontario, Canada) to enable spectral focusing based hsSRS microscopy (23, 24). In summary, chirping the two ultrafast laser pulses though high index glass from ~200 fs to about ~2.5 ps allows for tuning of the relative temporal delay between the ultrafast laser pulses at the sample to variably evoke SRS resonances. The result is improved spectral resolution (~30 cm^−1^) compared to using transform-limited 200 fs pulses (~300 cm^−1^) without being limited by laser wavelength tuning speed. The result is a video rate nonlinear microscopy platform with 800 nm spatial resolution and approximately 30 cm^−1^ spectral resolution. After chirping, the beams were directed to a pair of scanning galvanometric mirrors. The face of the first scanning mirror was relayed to the back focal plane of a physiological imaging objective (Olympus XLUMPLN 20X 1.0 NA, water dipping) through a 4X magnifying 4-f imaging relay (Thorlabs SL50-2P and TL200-2P).

**Figure 1:**
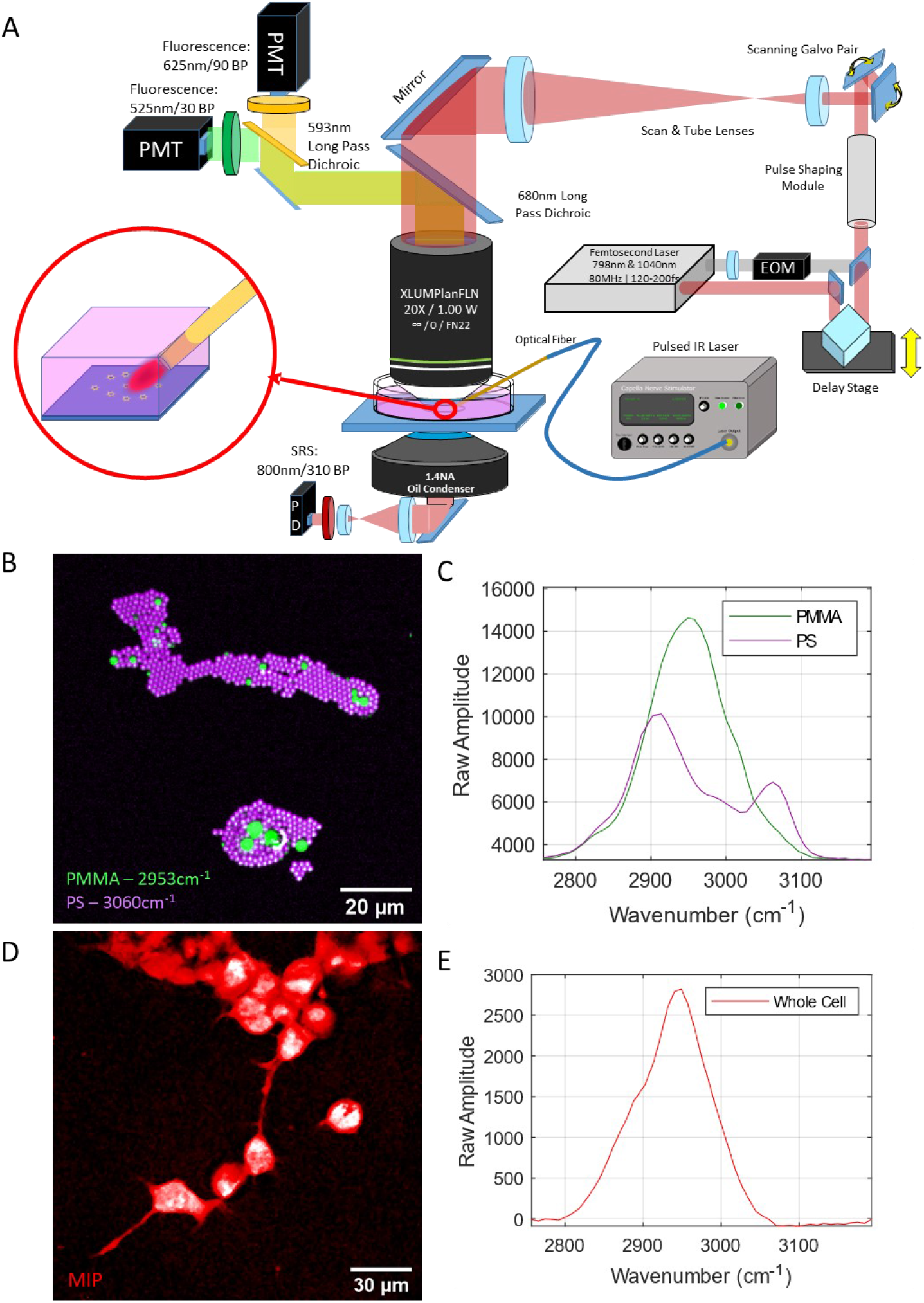
Experimental setup for SRS and fluorescence imaging of samples during IR exposure. (A) Imaging system schematic, (B) Standard poly(methyl methacrylate) and polystyrene (PMMA | PS) monolayer demonstrating spatial and (C) spectral performance of imaging system. (D) Maximum-intensity projection of the hyperspectral SRS image stack of live NG108 cells alongside their respective (E) whole-cell SRS spectra.

Detection for SRS – specifically stimulated Raman loss – was collected via transmission by a high-NA condenser lens (1.4 NA Oil, Nikon Instruments) directing light to a reverse-biased photodiode (APE, Gmbh., Berlin) behind an 850 nm centered, 310nm bandwidth optical bandpass filter (Semrock, Brattleboro, VT) to isolate the 798 nm laser line. The detected signal was subsequently demodulated with a lock-in amplifier (APE, Gmbh. Berlin) synced against the 20 MHz sinusoid signal driving the 1040 nm beam modulation. Any 20 MHz modulation transfer from the 1040 nm beam to the 798 nm beam was assumed to be attributed to stimulated Raman contrast. The temporal delay between the chirped 798 nm and 1040 nm laser pulses arriving at the sample was carefully tuned by varying the optical path length of the 798 nm laser beam with an optical delay stage (BB201, Thorlabs, Newton, NJ, USA). The relative delay between laser pulses over a span of 0.5 mm (or 1.6 ps) allowed for scanning of SRS resonance contrast over approximately 300 cm^−1^ between 2800 and 3100 cm^−1^. Additionally, this system also allows for for multiphoton fluorescence microscopy, which can be measured from epidetected light reflected from a 680 nm long pass dichroic mirror (Semrock, Brattleboro, VT) behind the objective lens in a non-descanned configuration. Bandpass filters for multiphoton fluorescence microscopy were selected to collect the green (525 nm center, 50nm passband) and red (625 nm center, 90 nm passband) emission profiles of the lipophilic dye di-4-ANNEPS. Images were acquired in a bidirectional point-scanning configuration. High-speed hsSRS imaging experiments were acquired with a 96×64 px sampling profile with varying pixel sampling densities between 1 and 4 µm/px. Imaging with 2-5 µs pixel dwell time and bidirectional scanning amounts to an effective imaging framerate approaching 150 Hz. Ultrafast laser average powers at the sample plane were measured to be about 10 mW for the 798 nm laser, and 25 mW for the 1040 nm laser – corresponding to 199W/cm^2^ and 497W/cm^2^ irradiances at each pixel, respectively. Assuming a constant 5 μs pixel dwell time, radiant exposures per pixel amount to 1.00 mJ/cm^2^ and 2.49 mJ/cm^2^. The substantial increase in power necessary for imaging in this configuration are due to the decrease in peak power within each ultrafast laser pulse during the chirping stage. Live cell viability was verified with all relevant imaging conditions by cellular uptake of propidium iodide (PI) and is described in *Live Cell Hyperspectral SRS Imaging*.

### hsSRS Spectral Focusing Calibration

A monolayer preparation of mixed polymer beads were used to calibrate the optical delay between pump and probe laser pulses as a function of SRS resonance. A mixed sample of poly(methyl methacrylate) – PMMA 1-10 μm diameter– and polystyrene – PS, 2 μm diameter – microspheres (PolySciences Ltd., Warrington, PA, USA) were diluted to a concentration of 0.002% w/v (each) in a solution of methanol (Fisherbrand, St. Louis, MO, USA). After mixing, 10 µL of diluted microbead solution were spread onto a #1.5 glass coverslip and left to evaporate for 25 minutes at room temperature. Once dried, samples were mounted dry onto a standard microscope slide and used for spectral calibration of hyperspectral SRS system by spectral focusing.

To calibrate the vibrational spectral dimension of hyperspectral imaging space, 50 sequential images were acquired of mounted polymer bead monolayers. Between each acquired image, the optical path length delay of the 798 nm laser line was stepped by 10 µm between each image, over a total of 500 µm or 1.6 ps of total optical path length delay. The peak SRS signal for the 2950 cm^−1^ resonance of PMMA was centered in the spectral scanning range to ensure sufficient spectral sampling. Manual segmentation of PMMA and PS beads from spectral stacks were performed and averaged across each spectral frame to provide high-fidelity spectra for both polymers. The known vibrational peaks of PS (2910, 3060 cm^−1^) and PMMA (2950 cm^−1^) were used as spectral fiducials (**Figure 1B** and **C**) to linearly interpolate a relationship between optical path delay of the chirped 798 nm laser pulse and the excited vibrational resonant mode. Calibrations were performed at the beginning of each day’s experiments to ensure spectral accuracy. The spectral resolution was observed to be approximately 30 cm^−1^.

### Infrared Neural Stimulation

Neural stimulation was performed by placing a bare 400 µm-diameter core low-OH optical fiber (Ocean Optics, FL, USA) in close proximity to samples (~450 µm) at a 30-degree approach angle into the sample plane of the microscope’s field of view (**Figure S1**). The optical fiber used for stimulation is connected to a pulsed laser diode centered at 1875 nm (Capella Nerve Stimulator, Aculight – Lockheed-Martin, Bothel, WA, USA). During imaging experiments, samples were exposed to a pulse train of 188 pulses distributed evenly over 1500 ms). Pulses were 400 µs in duration and were delivered at a repetition rate of 125 Hz. Radiant exposures on samples were varied by adjusting the peak current delivered to the laser diode, holding all dosing and geometric configurations constant. Radiant exposure calculations for stimulation were approximated based on power measurements performed externally in air and employing Beer’s law under the assumption of an absorption-dominated photon distribution – described in **Figure S1** and **Figure S2**. Infrared exposure levels for INS were selected based on their ability to elicit dynamic calcium responses (>2% increase, dF/F) in NG108 cells loaded with a calcium dye (Fluo-4-AM at 1 μM, ThermoFisher, St. Louis, MO, USA). Radiant exposures for no stimulation, sub-threshold, and threshold levels of stimulation used 0, 5.02, and 10.63 J/cm^2^, respectively.

### Phospholipid Multilamellar Vesicle Preparation

Multilamellar vesicles were used to obtain lipid-derived SRS spectra free of protein and nucleic acids signal in a biomimetic context. Multilamellar vesicles were prepared according to protocols provided from the supplier (Avanti Polar Lipids, Alabaster, AL, USA). Phosphatidylcholine (PC) derived from porcine brain tissue arrived dissolved in chloroform at a concentration of 2.5 mg/mL. The chloroform was evaporated from the lipid mixture with a stream of dry nitrogen overnight and mechanically resolubilized in phosphate-buffered saline solution at a concentration of 1 mg/mL. Vesicle mixtures were stored at 4°C and imaged within three days of preparation. Imaging was performed at room temperature. Size distribution of the lipid vesicle preparation was verified via dynamic light scattering to contain 1 and 5 μm diameter vesicles (Malvern Panalytical, Malvern, UK). MLVs were identified as multilayered spherical structures with SRS contrast tuned to 2910cm^−1^ (**Figure S4A**).

### Live Cell Hyperspectral SRS Imaging

Live cell imaging experiments of endogenous vibrational contrast with hsSRS were conducted with adherent cell preparations imaged in a physiologically balanced saline solution. Following placement of the fiber and calibration of the spectral axis against the known vibrational peaks of PS and PMMA beads, baseline hyperspectral image stacks were acquired for live cell samples. All images were acquired in a point-scanning approach with a 5 µs pixel dwell time and a spatial sampling density of ~500 nm/px. To improve signal to noise ratio of higher fidelity images, square fields of view between 320 and 512 pixels in size were acquired and 6 to 10 images were averaged together for each spectral position. For hyperspectral image stacks acquisitions, 50 images were acquired at evenly spaced intervals (10 μm) over 500 μm of optical path length delay – corresponding to a spectral range spanning approximately 2800 to 3100 cm^−1^. The resultant spectral image stack was taken as ground-truth cellular spectra to compare high speed imaging spectra of the cells during INS in subsequent experiments.

For high-speed imaging during INS on NG108 cells, as well as control samples of multilamellar vesicles and BSA solution, 5 µs pixel dwell time were employed to obtain imaging fields 96×64 pixels in size with a sampling density between 1.5 and 4 µm per pixel - enabling framerates of 33.4 Hz. For each of the 50 spectral position, cells were imaged continuously for 5 seconds, during which a train of stimulating infrared pulses is delivered at the first second of the imaging timeframe. Image acquisition and IR stimulation was coordinated through a customized TTL-triggering protocol with an external signal digitizer (Digidata 1550B, Molecular Devices, Sunnyvale, CA, USA). The ultrafast excitation laser is observed to be defocused at the sample plane due to the thermal gradient induced by the stimulating infrared laser (**Figure 2A**) was observable in each imaging timeseries as an exponential decrease, and subsequent return to baseline (**Figure 2B&C**), of nonlinear signal on imaging photodetectors. The shift in focal length as a function of laser power was calibrated using microbead (PMMA and PS) preparations and accounted for prior to each IR-stimulation trial on cells. The defocusing phenomenon allowed for precise temporal synchronization of time series across each spectral channel. After repeating and temporally aligning simultaneous imaging and stimulation time courses on lives cells for each SRS spectral position (n = 50), the temporal evolution of live cell endogenous vibrational spectra could be observed as a function of irradiation time and deposited energy. For spectral evaluation, the final ten sampling time points within the of IR exposure were averaged and reported – which was found to help reduce high frequency spectral noise to draw conclusions from. Spectra from stimulation experiments were pooled from n = 24 cells across ten different individual experiments of IR exposure. Each cell spectrum was normalized with respect to its integrated spectral intensity, and standard deviation of the spectra across all cells in each stimulation condition were calculated. The ‘no stimulation’ conditions are obtained from initial SRS signal from cells prior to each round of IR exposure and pooled from all stimulation conditions being compared. The shape of SRS spectra acquired at high frame rates (**Figure 3B**) were not found to noticeably differ from higher fidelity spectra (**Figure 1D**).

**Figure 2:**
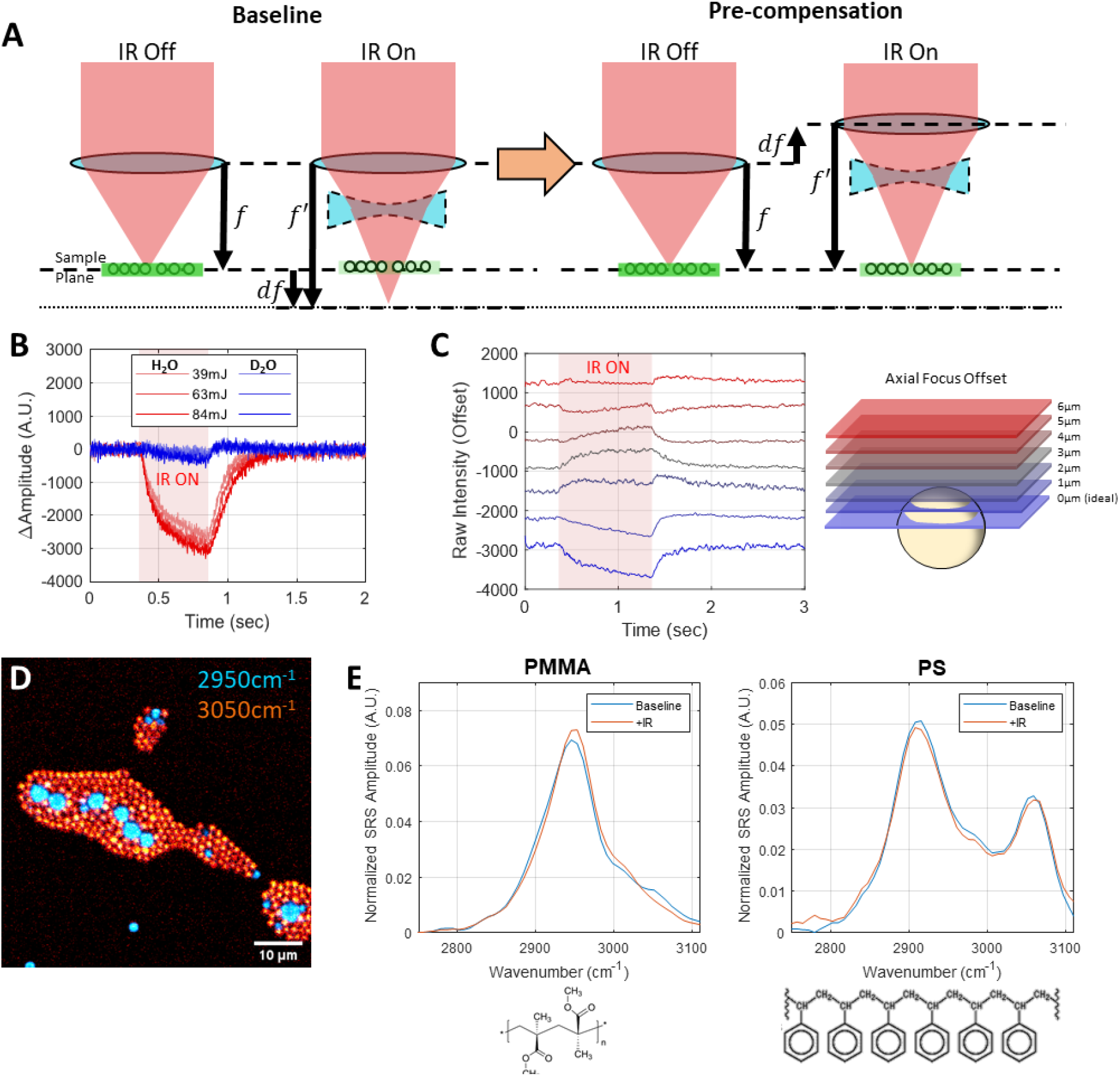
Explanation of defocusing phenomenon and the proposed experimental approach to circumvent it. (A) By adjusting the microscope focal plane to accommodate focal shifts induced by pulsed-IR neurostimulation within the microscope’s field of view, it is possible to recover some lost nonlinear signal due to defocusing. (B) The thermal gradient and subsequent defocusing artifact generated by INS in the microscope’s field of view is due to water absorption of INS light. Replacing H_2_O immersion with D_2_O immersion for imaging demonstrates that absorption of IR light is the driving force behind defocusing and signal loss. (C) Pre-compensating for INS-induced defocus by adjusting the focal plane position relative to our sample allows for nonlinear signal during INS. (D-F) Extrapolating this experimental approach across the wavenumber regions of interest allows for reconstruction of vibrational spectral dynamics during fast biophysical thermal events such as INS. **D)** Composite SRS image of PMMA and PS beads at 2950 and 3050 cm^−1^, respectively. Baseline and IR-stimulated spectra for E) PMMA, and F) PS reconstructed using the focus pre-compensation approach, with respective chemical structures for reference.

**Figure 3:**
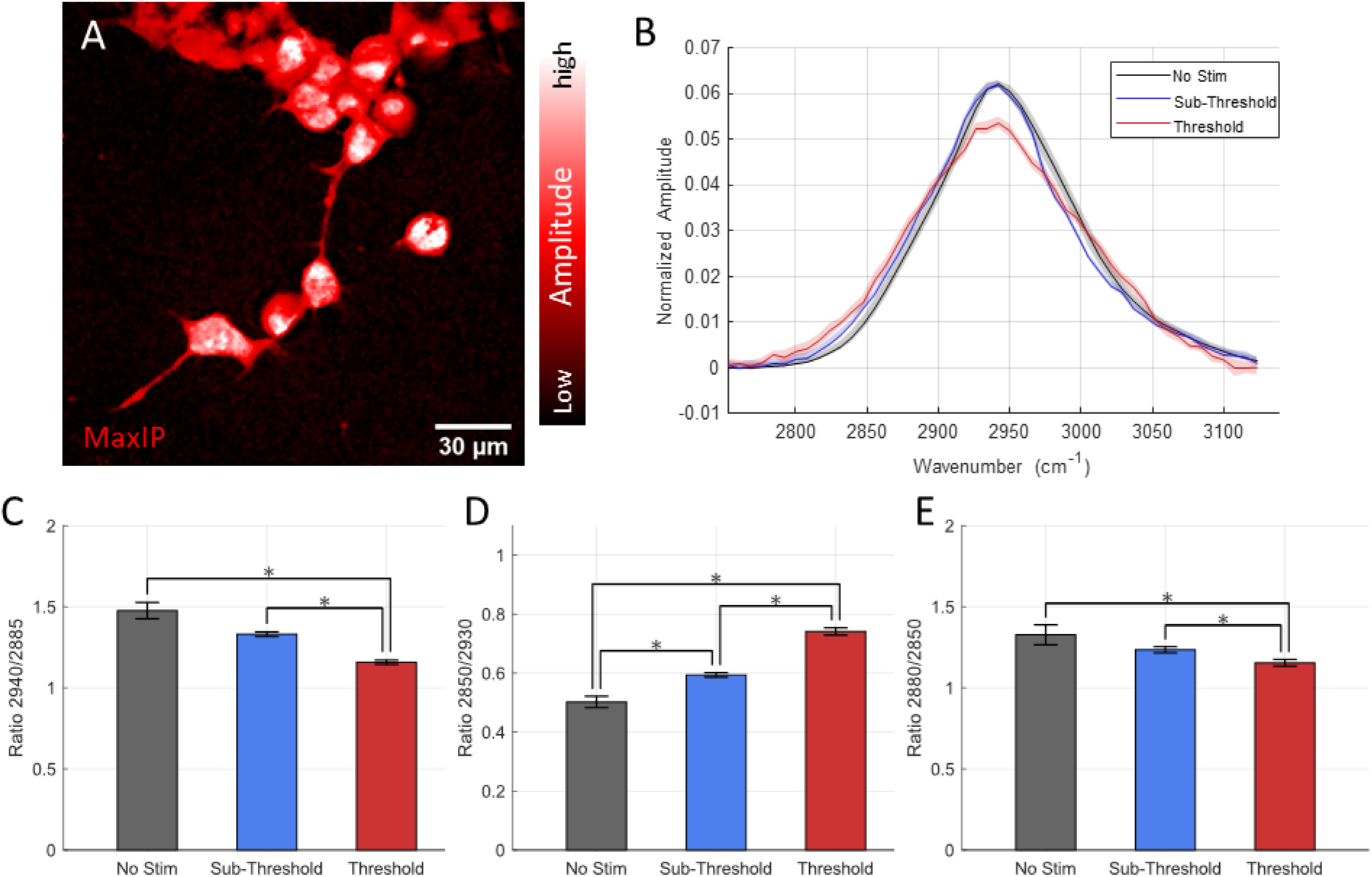
Vibrational Spectroscopic Imaging of NG108 Cells during infrared neural stimulation: (A)Maximum Intensity projection of NG108 spectral image stack from 2800-3150 cm^−1^ [n = 50 images]. (B)Average SRS spectra obtained from NG108 cells during infrared neural stimulation of at and above activation threshold radiant exposures [n = 10-24 cells per group]. Peak ratio comparisons indicative (C) asCH_2_/asCH_3_ as a measure of trans-to-gauche isomerization of lipid tail groups, (D) symCH_2_/symCH_3_ as a measure of increased polar headgroup association with water due to membrane packing order decrease, and (E) asCH_2_/symCH_2_ as an indicator of decreasing acyl chain packing order. *indicates p < 0.05

To verify cell viability during IR exposure, NG108 cells were subject to the hsSRS and stimulation protocol described above while simultaneously monitoring for cell damage via positive fluorescence staining of cell nuclei with propidium iodide. Imaging protocols were kept identical as previously described while supplementing the cell imaging medium with 1 µM propidium iodide (Thermo-Fisher, Natick, MA, USA). Cell morphology was additionally monitored throughout the experiment by comparing high fidelity images (< 1 μm/px sampling density) of the cells before and after imaging at their peak SRS resonance contrast at 2930 cm^−1^.

### di-4-ANNEPS Ratiometric Fluorescence Imaging

Imaging protocols were adapted from previously published work (25). Briefly, a loading solution of 4-(2-(6-(Dibutylamino)-2-naphthalenyl)ethenyl)-1-(3-sulfopropyl)pyridinium hydroxide (or di-4-ANNEPS) was prepared by diluting an aliquot of 4mM stock solution in dimethyl sulfoxide (DMSO) in neurophysiological saline to a final loading concentration of 2 µM. NG108 cells were incubated in the dark at 37°C, 5% CO_2_ and 95% relative humidity for 25 minutes, before being rinsed and maintained in fresh neurophysiological saline solution free of dye for fluorescence imaging. To image di-4-ANNEPS fluorescence, two photomultiplier tubes (PMTs) configured for non-descanned epifluorescence detection. Fluorescence emission was split by a 593nm long pass dichroic mirror and subsequently filtered with either a 525nm/25, or 625nm/45 optical bandpass filter before reaching PMT detectors (Semrock, Brattleboro, VT, USA). Ultrafast laser excitation for multiphoton fluorescence was tuned to 960 nm to optimally excite di-4-ANNEPS. For high-speed imaging, images were acquired as 96×64 pixel images between 0.5-4.0 μm/pixel sampling densities with 5 μs pixel dwell times to yield 33.4 Hz framerates. Excitation laser intensity for imaging was maintained below 10 mW at the sample plane. The SF11 glass rods used to chirp the laser pulses for hsSRS imaging were removed for ratiometric fluorescence imaging, resulting in ultrafast laser pulse width approaching 200 fs at approximately 80 MHz.

During a 5-second imaging period, stimulating IR light was delivered to di-4-ANNEPS stained NG108 cells via a 400 µm core multimode optical fiber immediately adjacent to the microscope’s field of view. Varying levels of radiant exposure were delivered to cells (0-44 J/cm^2^) and the resulting fluorescence intensity changes were compared across stimulation conditions. Calculations for conventional polarization, as well as a modified version of general polarization (**Figure S6**), were derived to compare conventional assessments of lipid packing with that observed with hsSRS.

### Data Processing, Analysis, and Visualization

#### Hyperspectral SRS Imaging Data

Raw data acquired from the imaging experiments were collated and sorted into multidimensional stacks of 16-bit TIFF stacks separated by time and wavenumber using a customize processing pipeline in Fiji leveraging the Bioformats plugin (26, 27). Average intensity projections of multidimensional (spectral, temporal) image stacks in time and spectral space were used to generate a mask to segment cells geometries. A general region of interest identified from the resultant masks were applied to the raw multidimensional image stack to extract spectral and temporal data from features of interest (e.g. beads, cells). To segment individual cells, a 2-pixel Gaussian blur was applied to the average intensity projection of the multidimensional image stack and contrast local histogram equalization was performed to reduce cell signal intensity variations between cells. Post-hoc flat field correction of imaging field heterogeneity of images was implemented by scaling pixel intensities relative to the average intensity projection gaussian blurred with a kernel equal to 0.25-0.5x the largest dimension of a particular image. Prominent peak locations in the image are identified. The filtered average intensity projection is subsequently segmented via Otsu segmentation. The resulting mask and previously identified peak locations are fed into an seeded watershed segmentation algorithm which reliably separates and segments induvial cells as their own ROIs with minimal cell-to-cell overlap (28). Edge maps of cells were acquired by subtracting the watershed-segmented mask from itself following an erosion operation, which reliably identifies borders in a cell-specific manner. The resultant regions of interest are applied to the raw stacks to extract the mean amplitude, standard deviation of signal or amplitude measurements, and their respective centroid locations in image-space for each spatial and temporal point. This process is automated as a macro procedure in FIJI and is freely available with raw data examples as supplementary information. Depicted images provided in the manuscript are derived either from single frames at specific wavenumbers of interest or maximum intensity projections of spectral image stacks. For visualization purposes in publication, intensity scaling for all images were adjusted linearly.

All hsSRS spectra are smoothed with a 3-point sliding Gaussian window and normalized with respect to their integrated spectral area. Since the intent of the study is to compare the relative spectral shapes of each sample, an integrated spectral normalization was chosen to facilitate this interpretation. Error associated with each plot is presented as standard deviation of all averaged spectra obtained for a given experimental trial. Each individual bead was taken as one sample, and different trials were taken as independent observations for statistical analysis purposes. For peak ratio comparisons, vibrational resonance intensities were calculated utilizing a cubic spline interpolation of the measured spectral data and it’s respective standard deviation. Comparisons of peak ratios were assessed using a student’s 2-sided t-test, where errors associated with ratiometric comparisons were calculated based on the propagation of error of the interpolated standard deviations (statistical significance was denoted by * for *p* < 0.05, ** for *p* < 0.01). All quantitative work was performed in MATLAB (Mathworks, Natick, MA, USA) using native functions. All bar graphs were created using the superbar package.

#### Ratiometric Fluorescence Analysis of di-4-ANNEPS Data

Processing of ratiometric fluorescence data is derived in part from previous work (29). Raw image stacks of green (lipid membrane gel phase - ordered) and red (lipid membrane liquid phase - disordered) spectral emission channels are acquired simultaneously at a 33.4 Hz framerate. Conventional general polarization (GP_conv_) was calculated using the following equation (29): Gwas calculated using the following equation (ref)

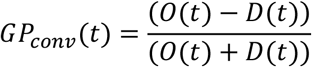

The raw image intensity differences between the green (ordered, O(t)) and red (disordered, D(t)) imaging channels were divided by the sum of both channels for each timepoint in the image stack for each experiment. Decreases in GP_conv_ value generally suggest decreases in membrane packing order. Average GP values as a function of time were calculated and each cell’s GP value was taken as an average GP of all pixels contained in each cell’s ROI. Cell segmentation similar to those segmented for SRS images utilizing a seeded watershed method was performed. However, since di-4-ANNEPS labels the extracellular membrane preferentially, a Huang threshold mask of raw disordered spectral fluorescence intensity images were obtained to determine cell boundaries and a binary fill operation was employed to identify areas in the image that contained cells. The lack of lipid-stained fluorescence in cell nuclei was used to identify center points of cells. The raw disordered fluorescence channel image was smoothed with a 2-pixel Gaussian filter and local minima in the images were used to approximately localize cell center points. These cell center points, as well as the cell position mask, and a distance map calculated from the cell position mask were fed into a seeded watershed algorithm in FIJI to yield segmentation maps of individual cells in a given experiment (26, 28). The regions of interest derived from the segmentation were subsequently applied to each imaging experiment, where time series of both raw fluorescence channels were obtained per cell and the resultant data was exported for processing and analysis in MATLAB (Mathworks, Natick, MA, USA). Statistical comparison of GP values across stimulation conditions was performed using a 2-sided student’s t-test and the magnitudes and standard error of means across the GP values were calculated across all individual cells in a particular experimental condition (statistical significance denoted as * for p < 0.05).

For image visualization, adapted from previous work (29), 8-bit depth raw fluorescence intensity images from the disordered fluorescence channel were multiplied by each color channel of an color red-green-blue (RGB) format image representing the calculated GP images with the desired false-colored look-up table of preference. The resulting images yield an image where pixel brightness represents intensity and color represents calculated conventional general polarization – which are used purely for visualization purposes. All rescaling of intensities in images are linear and performed for clarity of cellular morphologies and biophysical properties in print (**Figure 4A**).

**Figure 4:**
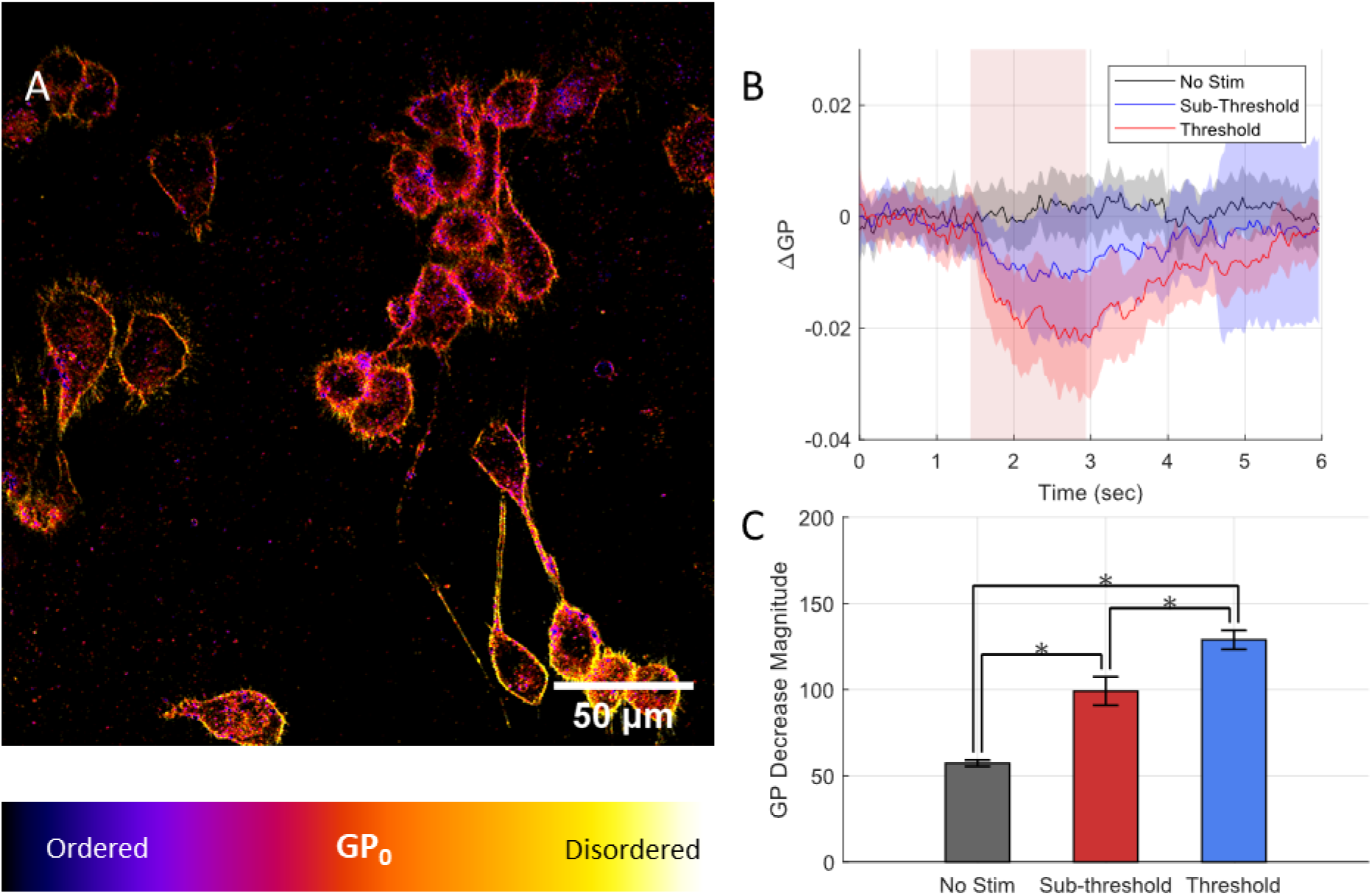
Relative changes in general polarization (GP) measurements in NG108 cells measuring dual-band fluorescence of di-4-ANNEPS verify changes in membrane order during INS. (A) Fluorescence intensity images overlaid with calculated initial GP values of NG108 cell cultures loaded with di-4-ANNEPS. (B) Relative changes in adapted general polarization metrics NG108 cells during varied doses of IR stimulation. Decreases in relative general polarization are indicative of decreases in relative extracellular lipid membrane packing order which agree with hsSRS observations. Error traces represent standard deviation across all cell responses [n = 50-109 cells]. (C) Magnitude of GP decreases across sub-threshold [5.02J/cm2] and threshold [10.63 J/cm2] levels of radiant exposure. Error bars represent SEM across all cells within each condition. * indicates p < 0.05.

Due to large variations in total fluorescence measured in any given experiment due to thermal lensing during IR stimulation, the conventional method of calculating GP was found to be unreliable. Since we expect a decrease in overall fluorescence due to the decrease in effective collection efficiency during thermal lensing induced defocusing, the magnitude of changes in the denominator of the GP_conv_ equation are much larger than that of the changes in the numerator of the equation. To account for these effects, we developed an intensity invariant version of GP_conv_ to better reflect these dynamics mathematically over short experimental periods of time undergoing substantial changes in photon collection:

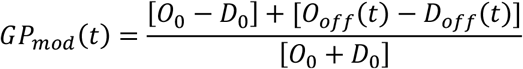

Where O_0_ represents initial ordered fluorescence levels, D_0_ represents initial disordered fluorescence levels

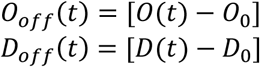

O_off_(t) represent the net change in ordered fluorescence relative to O_0_ as a function of time, and D_off_(t) represents the net change in disordered fluorescence as a function of time. O(t) and D(t) are the raw ordered and disordered fluorescence as a function of time, respectively. (**Figure S6B**). The alternative metric of modified GP (GP_mod_) emphasizes the raw difference in measured fluorescence intensity between the ordered and disordered fluorescence imaging channel without dividing by the sum of both image channels over time. Assuming the defocusing artifact between both channels results in an equal amount of defocusing and signal loss from each fluorescence channel, any changes in the relative difference between the fluorescence signals as a function of time is indicative of functional changes in lipid bilayer packing (**Figure S6**). For the purposes of this study, we are interested in determining the direction of GP changes – positive or negative - rather than it’s magnitude. This consideration makes GP_mod_ a convenient and applicable tool for our experimental approach.

## Results

### Thermal Lensing during IR Stimulation

Following confirmation of our instrument’s ability to obtain hsSRS image stacks from live NG108 cells (**Figure 1D** and **E**), initial experiments with IR stimulation during nonlinear microscopy (i.e. any coherent Raman modality, multiphoton fluorescence, or higher harmonic generation) resulted in a substantial loss in measured signal during IR exposure (**Figure 2B** and **C**) (22). This was apparent in both short periods of heating from a millisecond pulse of IR light (unpublished data), or pulse trains of multiple microsecond pulses of light. The shape of the disappearance and reappearance of the nonlinear signal appears to follow the shape of the expected heating and cooling dynamics that is typically observed during IR mediated heating (30)– suggesting that a temperature related phenomenon may be responsible for the loss in signal. Considering the goal of this work is to image the high-speed chemical dynamics in live cells during IR exposure, the loss of signal during this critical time period posed a challenge. To better understand the role of this signal loss with immersion medium temperature, a vegetable oil sample was imaged with SRS (2885 cm^−1^) through warmed immersion medium at a range of physiologically relevant temperatures. Temperature of the immersion medium was monitored by a thermocouple placed adjacent to the microscope’s field of view at the coverglass-immersion medium interface (**Figure S3**). Warmed deionized water (approximately 50 °C) was added between the objective and sample with the edge of vegetable oil sample placed in focus. Images were acquired continuously as the immersion medium slowly cooled to room temperature (22 °C). Contrary to the signal decrease observed during rapid IR heating (**Figure 2B** and **C**), this experiment showed that changes in immersion medium temperature revealed a positive correlation with temperature and SRS signal of vegetable oil. This data suggested that changes in immersion medium temperature on its own was not sufficient to explain the decrease in nonlinear optical signal during IR heating.

The refractive index of the objective immersion medium, water (H_2_O), is negatively correlated with temperature (31). This concept suggests that the spatial thermal gradients generated by the IR absorption from IR stimulation would defocus the ultrafast laser driving nonlinear contrast and thus reduce observed nonlinear optical signal. To test this hypothesis, the immersion medium for the objective lens was replaced with heavy water (D_2_O), which has a five-fold lower absorption coefficient at 1875nm than deionized water with nearly identical refractive indices (2). If the thermal gradient causes the decrease in nonlinear signal observed in the sample, then reducing the immersion medium’s IR absorption properties should reduce the magnitude of the nonlinear signal decrease during stimulation. The results shown in Figure 2B validates this hypothesis (**Figure 2B**) suggesting that the thermal gradient from IR stimulation was defocusing the ultrafast laser source resulting in a decrease in nonlinear signal (**Figure 2A**).

Since water’s index of refraction is negatively correlated with temperature, the thermal gradient generated during IR stimulation in front of the stimulation fiber and within the microscope’s field of view behaves like a negative lens during imaging. Imaging out of focus samples during IR stimulation would bring samples into focus (**Figure 2A**). This hypothesis was found to be true for both nonlinear imaging and IR transillumination imaging. By moving the microscope’s focal plane above the sample by a few microns prior to IR exposure, the samples (polymer microbeads in this case) would come into focus (**Figure 2C**). This precompensation of defocus was applied repeatedly across numerous spectral channels to generate a time resolved hsSRS profile of samples during IR stimulation, similar to previously employed approaches with hsSRS and electrophysiology (18, 32). This approach was verified by measuring several control samples: PS/PMMA microbead monolayer mixtures, 10%w/v bovine serum albumin solution in PBS, and large multilamellar vesicles of neurologically derived phosphatidylcholine (PC) and phosphatidylethanolamine (PE) in physiologically balanced neural saline solution before conducting experiments using live cellular samples.

### Verifying pre-compensation for thermal defocusing during hsSRS

**Figure 2D** shows a representative image of mixed microbead monolayers, highlighting PMMA in cyan using the band at 2950 cm^−1^(terminal methyl C-H resonance) and PS in orange using the band at 3050 cm^−1^ (aromatic C-H stretch resonance). The mixed bead sample was exposed to ~12 J/cm^2^ IR stimulation and the resultant spectra for both bead types are shown in **Figure 2E&F**. Relevant spectral band assignments for polymer microbead samples are summarized in **Table 1**. Infrared-exposed PMMA beads exhibit several distinct spectral changes upon heating – decreases in the 2880 and 2910 cm^−1^ resonances of skeletal C-H stretching, as well as relative increases in resonances at 3000 cm^−1^ and decreases at 3050 cm^−1^. Shifts in PS hsSRS spectra during IR exposure show relative increased vibrational activity around 2850 cm^−1^, implying the possibility of relaxed steric hinderance of skeletal sp^3^ CH_2_ symmetric stretching modes, while broadening the 3050 cm^−1^ peak attributable to aromatic sp^2^ C-H stretching and suggesting reduced steric hinderance around aromatic side chains. These observations show that utilizing a time resolved approach to obtaining hsSRS spectra of samples heated by pulsed IR light is feasible in highly Raman active idealized chemical samples.

**Table 1:**
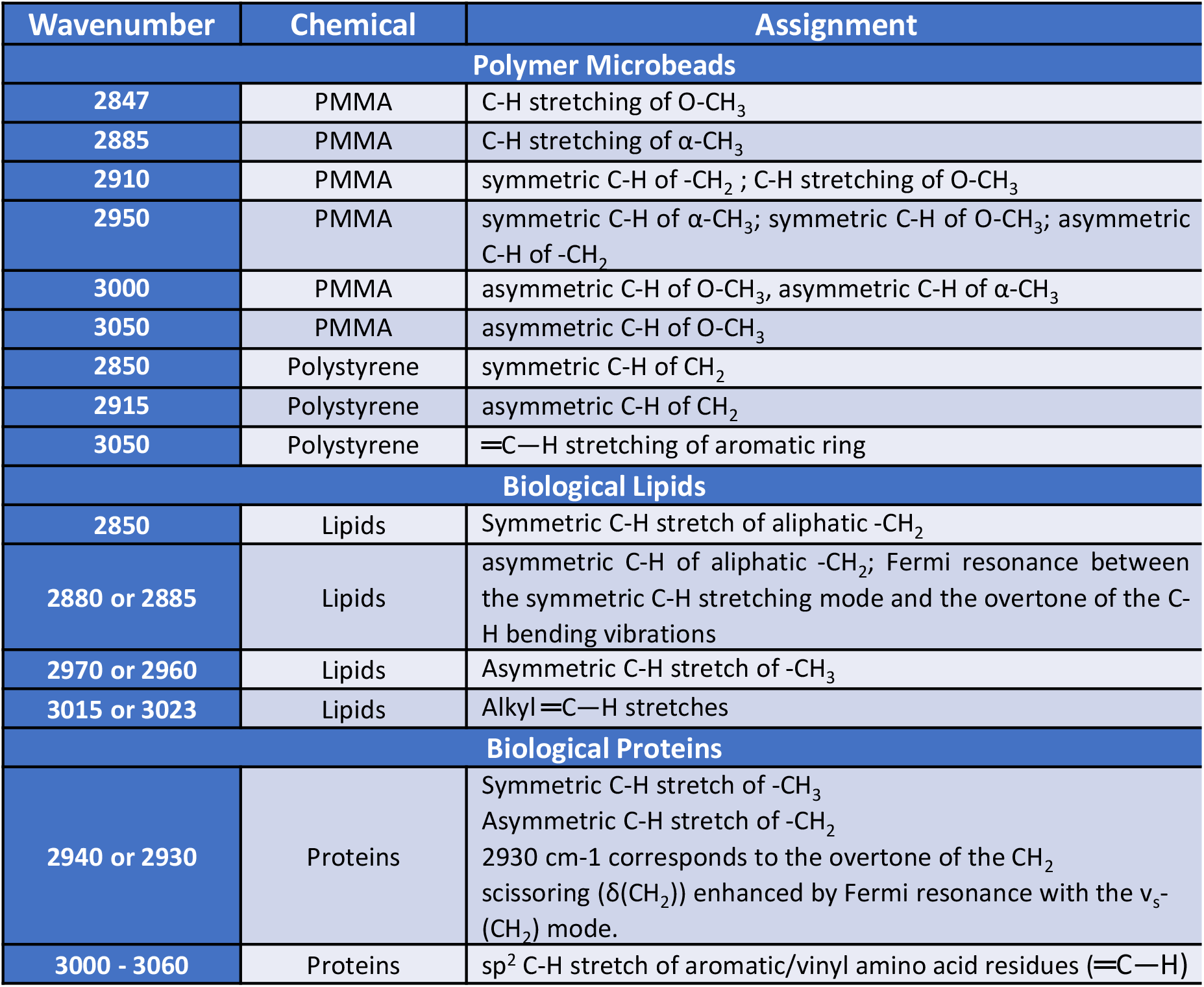
Raman spectral band assignments in the CH stretch region for control (23, 73, 74) and cellular (36, 37, 43, 75) samples

The dominant Raman scatterers in the 2800-3100 cm^−1^ spectral region primarily include lipids and proteins – with some marginal nucleic acid contribution (33, 34). Spatially and spectrally, nucleic acids are easy to separate in cellular images (35). However, since proteins and lipids in cells do not appear as spatially distinct as the resolution of our microscope, their distinct spectral information must be used to draw conclusions about their molecular dynamics. Understanding how proteins and lipids are separately affected by IR stimulation provides insight to the spectral shifts can be attributed to each biomolecule during live cell imaging. hsSRS imaging with IR stimulation was performed on separate aqueous preparations of biomimetic multi-lamellar vesicles (phosphatidylcholine - PC, neurologically derived, porcine sourced, Avanti Polar Lipids, Alabaster, AL, USA) and bovine serum albumin (BSA, 10%w/v) solutions.

An emulsion of multilamellar vesicles (MLVs) were imaged with hsSRS and focus precompensation during radiant exposures equivalent to threshold levels (10.63 J/cm^2^) of IR exposures in live cells. These vesicles serve as a coarse chemical representation of cells to provide an isolated lipid preparation, free of protein or carbohydrate contribution to vibrational spectra. Infrared-exposed MLV spectra (**Figure S4A, B**) show distinct shifts in lipid molecule resonances relevant to lipid molecular packing order. Relevant spectral band assignments for biological lipid samples are summarized in **Table 1**. The 2850 cm^−1^ symmetric aliphatic C-H stretch resonance is markedly decreased, along with its Fermi resonance at 2880 cm^−1^. Meanwhile, sp^2^ C-H stretching resonances associated with unsaturated aliphatic chain motifs at 3010 cm^−1^ are substantially decreased. Crucially, ratiometric comparison of 2880 and 2850 cm^−1^ shows reduced rotational restriction in alkane chains, or a decrease in aliphatic tail packing order within the hydrophobic region of the membrane (**Figure S4C**). This is further supported by the observed decrease in the ratio of 2940 to 2830 cm^−1^, which relates to increases in the solvent interaction with lipids (**Figure S4C**). These observations suggest that thermodynamic changes in lipid vibrational signatures during IR stimulation are discernable with hsSRS.

To characterize protein vibrational signature changes during IR-induced heating, the edge of a 10%w/v BSA solution meniscus was imaged with hsSRS using radiant exposures equivalent to threshold levels of IR exposures in live cells (**Figure S4D**). Changes in protein spectra during IR exposure appear to be negligible (**Figure S4E**). Furthermore, the contribution of protein vibrational spectra in ratiometric comparisons that reveal significant changes in MLV samples appear to contribute negligibly to IR-exposed changes in the BSA sample (**Figure S4F**). It is worth noting that the amino acid constituents of BSA, a water-soluble protein, may not be directly representative of a transmembrane protein one would observe as a component of the extracellular membrane or intracellular organelles. However, the data supports previous work showing that the shape of protein spectra in the CH-band region of the Raman spectrum do not appreciably change with temperature (36, 37).

### hsSRS of Neural Cell Models during INS

With the spectral changes in biomimetic samples established, hsSRS imaging during IR stimulation was conducted in an *in vitro* neural cell model - a neuroma-glioblastoma hybridoma cell line (NG-108-15, Sigma-Aldrich, MO, USA). The NG-108 cell line was used as a practically robust and experimentally resilient neuronal cell model for hsSRS imaging. These cells are an accepted electrodynamic model of *in vitro* neurons and have been used in the past successfully to study electrodynamics evoked by IR stimulation (19, 20, 38). **Figure 3A** shows a maximum intensity projection of an hsSRS spectral image stack to highlight the morphology of NG-108 cells. Successful stimulation with pulsed IR light were verified in separate experiments (unpublished) of NG108 cells loaded with a calcium-sensitive dye, Fluo-4-AM at 1µM in balanced saline for 45 minutes. Two-photon fluorescence and SRS centered at 2880 cm^−1^an asymmetric sp^3^ CH_2_ resonance dominantly from lipids –images were acquired simultaneously during IR stimulation of NG108s at a range of IR doses until noticeable increases in calcium-dependent fluorescence responses were evoked (>2% increase in dF/F). Levels of IR evoking consistent intracellular calcium responses across the microscope’s field of view were referred to as threshold levels of exposure. Cells were imaged with hsSRS during IR stimulation with threshold and subthreshold (about half of threshold levels) doses of IR light.

The resultant area-normalized hsSRS spectra of NG108 cells under baseline (unstimulated), subthreshold, and threshold stimulation conditions are shown in **Figure 3B**. Relevant spectral band assignments for biological samples in the CH-stretch region are summarized in **Table 1**. Shoulders appearing at 2850 cm-1 during stimulation are indicative of relatively increased vibrational resonant activity from symmetric aliphatic C-H stretching in lipid tail chains. Decreases in the relative intensity ratio between 2940 and 2885 cm^−1^ (**Figure 3C**) suggest a decrease in packing order within the hydrocarbon tails of the lipid molecules due to *trans*-*gauche* isomerization of sp^3^ hydrocarbon chains. Interestingly, the 2850 cm^−1^ shoulder appears to increase in spectral intensity relative to the associated Fermi resonance at 2880 cm^−1^, possibly suggesting a reduction of intermolecular steric hindrance between aliphatic lipid tails, or more rotational freedom of hydrocarbon chains. These observations were quantified by calculating the intensity ratio between 2850 and 2940 cm^−1^ (**Figure 3D**), as well as 2880 and 2850 cm^−1^ (**Figure 3E**). These metrics respectively offer a quantification of lipid tail chain packing order which was previously hypothesized to decrease during IR stimulation (3). **Figure 3C-E** shows these intensity ratios from NG-108 whole cell spectra obtained at baseline, sub-threshold, and threshold levels of INS previously established to elicit calcium transients. Statistically significant differences (p < 0.05, indicated with asterisk) in these ratios suggest decreased hydrocarbon tail chain packing in cellular lipid membranes. Notably, in each comparison, the ratios calculated for subthreshold exposure fall between unstimulated and stimulated conditions. Of particular note, the shoulder around 3030 cm^−1^ – which is a sp^2^ CH (methylene) resonance assignable to CH bonds at points of unsaturation in lipid hydrocarbon tails – appears at the threshold stimulation but is reduced in the subthreshold and no stimulation cases (**Figure 3B**).

The hsSRS spectral acquisition as described above requires cells to be exposed to 50 different rounds of IR stimulation – possibly damaging the cells and yielding biologically irrelevant observations. Though no morphological changes were observed in the stimulation experiments, cell viability was verified after repeated IR exposure. Exposed NG108 cells were imaged with multiphoton fluorescence to track the uptake of a cell damage indicator – propidium iodide (PI) – simultaneously with SRS tuned to the 2940 cm^−1^ CH_3_ resonance. Cells were imaged through 50 rounds of stimulation, using parameters similar to those used during a live cell hsSRS imaging experiment (

**Figure S5A**). Some cell swelling was observed morphologically, but no uptake of PI was observed (

**Figure S5B**)– suggesting that the repetitive nature of hsSRS acquisition did not have any immediate impact on acute cell viability.

### Ratiometric fluorescence imaging of functional lipid dye during INS verify changes in lipid bilayer packing order

Ratiometric fluorescence of di-4-ANNEPS emission, a probe of membrane packing order, was employed to verify cellular lipid dynamics as observed in vibrational spectra (25). Di-4-ANNEPS rotoisomerization is known to be dependent on fatty acid tail chain packing order in lipid membranes. During IR stimulation, if lipid tail chain packing order is decreased, a similar decrease in general polarization (GP) metric should follow. In place of the conventional approach for calculating GP, intensity-invariant adaptation of GP was utilized to circumvent the defocusing effect during IR stimulation (detailed in Methods and **Figure S6**). **Figure 4A** depicts an intensity image of di-4-ANNEPS loaded NG-108 cells overlaid with color denoting GP calculation at each pixel. **Figure 4B** and **C** show the mean single cell GP time traces and their standard deviations for each dosing condition. The intensity-invariant GP of di-4-ANNEPS (**Figure S6**, see Methods) shows substantial decrease in GP as a function of IR stimulation dosage (**Figure 4C**). A decrease in GP suggests a decrease in lipid chain packing order during IR stimulation supporting the hsSRS observations.

## Discussion

Our current understanding of label-free directed energy neuromodulation continues to raise questions about their mechanistic bases. An improved understanding of INS mechanisms provides a fundamental framework for the development of future innovative neuromodulation technologies. Here, we provide an approach that uses hsSRS microscopy to gain insight to the role of lipid dynamics in live neural cells during INS. Most traditional methods to observe lipid-specific dynamics (e.g. isolated lipid preparations, electrophysiology, x-ray diffraction, neutron scattering) in cells in real time suffer from lack of specificity or biological compatibility. Methods that utilize fluorescent tags (e.g. fluorescence correlation spectroscopy, fluorescence recovery after photobleaching) provide insight into the dynamics of lipids in live cells but are inherently indirect. The goal of this work was to directly observe the biophysical dynamics of INS with a vibrational spectroscopic approach in live neural cells. Using the intrinsic Raman contrast of lipids, spectroscopic insight would help clarify the mechanistic role of lipid dynamics in INS. Our demonstration of characterizing and precompensating for dynamic defocus during INS with hsSRS is a novel approach in biomedical microscopy that is applicable to studying the molecular biophysics of live cell models more generally.

Photothermal events are notoriously difficult to address with biological microscopy due to the relationship between temperature and refractive index in water. While bulk changes in sample temperature can impact optical aberrations in microscopes, spatial thermal gradients that vary on the order of the microscope’s field of view can have significant impacts on the refraction of light into the sample (**Figure 2B**). Accounting for defocusing actively on millisecond timescales may be possible with dynamic adaptive optics approaches but is far from trivial to implement. Instead, our approach to adjust for IR-induced defocusing of the fluorescence excitation empirically (**Figure 2A, C**) – though coarse compared to adaptive optics – enables us to gather useful insight to the biophysical phenomena associated with INS (**Figure 3**). The reliable timing of stimulation can be leveraged to employ a time-resolved spectroscopy approach to hsSRS imaging at high framerates. In doing so, we demonstrate that high-speed vibrational dynamics can be resolved in live cell preparations safely to yield biologically meaningful observations. In studying INS using high numerical aperture microscopy, where IR induced deflections in focal length can equal or exceed the depth of focus of a particular imaging objective, we urge others to interpret their results cautiously. Thermal defocusing can have a disproportionate impact on single-channel intensiometric-based measurements and need to be carefully accounted for (**Figure S6**). In cases where intensity noticeably changes during exposure, we encourage others to employ ratiometric or multi-spectral approaches to allow for defocusing artifacts to be readily accounted for. With fluorescence microscopy, where quantum yield, fluorescence intensity, and spectral profiles are well known to be sensitive to both heating and defocusing (39–41), having simultaneous or time-resolved multispectral reference bases will allow for such artifacts to be accounted for in post-processing.

There are several spectral changes in the CH-stretch region of the Raman spectrum (2800-3100 cm^−1^) that one might expect to see if the current INS mechanistic model was valid. *Trans*-*gauche* isomerization, or rotoisomerization, of sp^3^ hydrocarbon chains – primarily associated with lipid hydrophobic tail groups in Raman imaging – can give rise to a number of steric effects that drive lipid membrane deformations (36, 37, 42, 43). Specifically, lipid packing order – or the ability for lipid molecules to stack neatly alongside each other within the membrane leaflets – was hypothesized to decrease with elevated temperature during INS. Rotoisomerization in membrane lipids geometrically shortens acyl tail groups, resulting in membrane thinning. While quantifying the absolute deformation of lipid membrane thickness with SRS would require additional calibration experiments, relative indicators of molecular interactions can be quantified with hsSRS. An increased quantity of gauche rotamer within the hydrophobic region of the membrane leads to geometric acyl tail shortening and sterically drives lipid molecules apart from each other. The result is a decrease in membrane packing order. In the CH-stretch region of the Raman spectrum, relative changes in symmetric (2850 cm^−1^) and asymmetric (2880 cm^−1^) aliphatic C-H stretching indicate shifts in molecular packing order due to changes in the rotational freedom of hydrocarbon chains in lipid tails. Raman signal at these resonances is largely attributed to biological lipids (**Figure S4**) (33). A decrease in the ratio of 2880 and 2850 cm^−1^ during INS (**Figure 3E**) is indicative of a ‘loose’ packing order between lipid molecules or an increase in trans-gauche isomerization (44, 45). The rotoisomerization of lipid tails is well known to both decrease membrane thickness and increase the area of each lipid molecule’s solvent interactions (46–48). Changes in the ratio between 2940 and 2885 cm^−1^ offer insight to water interaction with lipid molecules, which should increase with temperature. The data show a decrease in the ratio between 2940 and 2885 cm^−1^ (**Figure 3C**), which is in line with the idea that lipid molecules expand within the membrane leaflets to leave room for more potential solvent interactions (e.g. hydrogen bonding) with elevated temperatures. The IR dose dependance of this observation further suggests that the relative degree of isomerization correlates with levels of IR exposure that would evoke neural activity *in vitro*. The observations of a progressive increase in isomerization with IR exposure support the existing mechanistic model of INS, where transient temperature changes are accompanied by changes in physical bilayer geometry.

The shoulder appearing around 2990 and 3030 cm^−1^ during INS in cells (**Figure 3B**) arise from relative increases in vinyl C-H resonances, which correspond to points of unsaturation in lipid tail acyl chains. Relative increases in vinyl C-H signal can arise from reduced steric hinderance of sp^2^ C-H stretching as well as compositional or membrane potential related changes when the lipid bilayer undergoes thermal changes. Curiously, the appearance of the 3030 cm^−1^ shoulder in threshold stimulated cells was reduced in sub-threshold levels of stimulation. This resonance at 3030 cm^−1^ may provide a key marker for neural biophysics during INS.

The vinyl C-H portion (2980-3100 cm^−1^) of the C-H stretch region does contain SRS signal contributions from proteins– particularly from amino acid residues such as tyrosine, phenylalanine, and tryptophan.

These amino acids play a key structural role in stabilizing hydrophobic domains of transmembrane proteins in the cell membrane. Control experiments observing the IR-related dependance of BSA SRS spectra in solution (**Figure S4**) as well as evidence from others (36, 37, 49) reinforce that thermally-mediated changes in protein dynamics are not major contributors in the CH stretch region of the Raman spectrum. As such, we conclude protein signal contributes minimally to the photothermal mediated SRS changes that would be expected during INS. Others have attributed relative decreases in 2930 cm^−1^ signal to changes in cellular membrane potential, enabling the visualization of neuronal action potentials with SRS microscopy (18, 32). These spectral changes were attributed to the decrease in positively-charge proteins electrostatically accumulating at the extracellular membrane surface when a cell is at its resting membrane potential. A reduction in membrane potential was suspected to reduce membrane-associated proteins in solution at the cell membrane surface. Our results show a considerable reduction in relative 2930-2940 cm^−1^ signal during INS (**Figure 3B**), thus electrostatic association of soluble proteins with cell surfaces may play some role in our results. Several experimental details suggest that membrane potential changes from electrostatic protein association would not be contributing to our spectra. Defocusing artifacts make it difficult to obtain conclusions about absolute molecular concentrations during INS (**Figure 2A-C**). Practically, our approach to region of interest selection, non-balanced detection, and imaging medium formulation confounds any comparability of our results with these previous studies.

However, Lee et al. did employ a similar time-resolved approach for acquiring SRS spectra as a function of membrane potential – demonstrating the utility of such an approach for certain types of experiments beyond photothermal phenomena.

The physical changes in the lipid bilayer during rapid heating with IR light are thought to give rise – at least in part – to the cell capacitance increase that drives cellular depolarization during INS (2, 3). Our results (**Figure 3**) support the idea that the lipid bilayer undergoes some thermally mediated chemo-physical change during INS that is observable via vibrational imaging and correlate with the level of delivered stimulus. While these findings are promising, they do not definitively support that bilayer deformation is directly causal to the stimulatory effect of INS. Though beyond the scope of this work, questions remain about how transmembrane ion channels may be independently sensitive to lipid membrane geometry and thermodynamics. Lipid thermodynamics are known to affect the conformational and functional properties of transmembrane ion channels (50–53). It is not clear whether the capacitive effect or the actual physical change in the lipid bilayers themselves give rise to stimulatory phenomenon. It is difficult to decouple chemo-physical and thermal electrodynamic changes in biologically relevant preparations. A preparation of lipid vesicles or cells expressing voltage gated ion channels loaded with a UV photo-switchable lipid analogue (e.g., containing an azobenzene moiety in the tail group) may be a useful experiment. The photo-switching property of such synthetic lipids would allow for optical control of membrane packing order with substantially reduced photothermal effects.

The current hypothesis for how INS occurs is that rapid heating causes a capacitive inward current that can depolarize neurons and lead to action potential generation (2). This capacitive current is thought to arise from biophysical changes within the extracellular membrane – namely *trans*-*gauche* isomerization of lipid acyl tail chains – that change the physical dimensions of the extracellular membrane due to temperature elevations (3). This deformation is accompanied by a movement of membrane-associated charge that – when hot and fast enough – can generate an inward current that depolarizes cells. The model of this phenomenon relies on steady-state chemical assessments of synthetic lipid bilayer geometry (54, 55). The changes in bilayer geometry are used to inform a computational electrodynamic model that is compared against previous experimental work (2, 38). While the model of chemo-physical and electrodynamic phenomena convincingly reproduces experimental data, capacitance changes and cellular electrodynamics are ultimately influenced by more than lipid dynamics alone *in vitro* and *in vivo*. Our work here provides direct evidence that lipid membrane dynamics are actively changing during INS in neural cells *in vitro*. The causality of this phenomena remains to be proven. But the insight provided by our work shows how lipid membrane dynamics can be leveraged to selectively modulate cellular physiology.

Our SRS spectral observations are supported by an additional gold standard means of measuring lipid dynamics in real-time – ratiometric fluorescence of a lipophilic dye, di-4-ANNEPS (**Figure 4, Figure S6**). The negative changes in GP during INS affirm the decrease in membrane packing order observed with hsSRS. The magnitude of the changes in GP scaled with the level of stimulus delivered (**Figure 4B** and **C**). The data further suggests that hsSRS can be leveraged as a complementary tool to study lipid biophysics alongside traditional fluorescence approaches. Others have applied hsSRS to observe lipid biophysics in synthetic preparations (16, 17), or to study lipid metabolism at the biomolecular level (56, 57). Stimulated Raman microscopy has not previously been applied to the study of biophysical thermodynamics at sub-second timescales. Our work explores a temporal regime of live cell biophysics that few have ventured into with SRS. This work provides a practical extension to the existing work around hsSRS development while shedding light on a question pertinent to the field of optical neuromodulation.

While the implementation of hsSRS here can resolve high speed spectral dynamics well below a second, it does take several minutes to build observations of events on a spectral basis. In situations where repeated perturbation of cells is not practical, the same approach can be implemented with a drastically reduced number of spectral channels. Alternatively, multispectral approaches leveraging simultaneous acquisition of multiple resonances would be advantageous. To account for the defocusing artifacts described here, at least two spectral channels need to be acquired to accurately draw conclusions – thus single-shot perturbations are not readily applicable with the demonstrated approach here. The fast rates of development in bioimaging with SRS show promise in pushing SRS based imaging methods to their limits. Our work shows that hsSRS can be applied to a range of lipid biophysics experiments as a complement to more conventional fluorescence-based approaches. In contrast to fluorescence-based approaches that rely on indirect readout from reporter molecules interacting with lipids in the cell membrane, vibrational contrast like that of hsSRS enables direct inference to be made specific to lipids at the intra- and intermolecular levels. As technology in coherent Raman imaging continues to improve with better lasers, detectors, and signal processing strategies, we can expect to see extensions of hsSRS to address many other areas of lipid biophysics and beyond. Currently, signal to noise limits the real-time performance of hsSRS in the fingerprint region of the Raman spectrum (400-1700 cm^−1^). In future studies we propose to study the fingerprint region which provides more information about other biomolecules, such as DNA, RNA, and carbohydrates, which can be used to study macromolecular phase separation phenomena, chromatin dynamics, or glycogen metabolism directly without exogenous labeling.

Furthermore, coherent Raman imaging can be readily performed simultaneously with other nonlinear microscopy modalities (22). Multiplexing modalities might enable studies into how lipid membrane biophysics can influence biological dynamics with conventionally accepted molecular reporters. With this in mind, hsSRS has promising potential for a diverse range of bioimaging applications.

Alternative approaches utilizing deuterated lipid preparations to shift lipid-specific resonances into the “silent window” of the Raman spectrum (1700-2700 cm^−1^) may offer additional insight into the role of vinyl C-D resonances in the biophysics of INS (12, 58–60). However, the applications of deuterated lipids may not be easily replicable in live cells as it can interfere with the hydrogen bonding dynamics crucial to cell membrane integrity. Currently, fast implementation of hsSRS is technically hampered by the signal-to-noise performance in the fingerprint window of the Raman spectrum (400-1700 cm^−1^).

Utilizing other features of the Raman spectrum that are more directly attributed to lipid tail chain rotoisomerization (e.g. the skeletal vibrational C-C modes between 1030 and 1150 cm^−1^, as well as C=C stretching modes around 1650 cm^−1^) might provide more direct mechanistic insight to INS once possible (44). Some promising newer spectroscopic and computational denoising methods that circumvent these noise issues are gaining popularity, but still require careful validation for high-speed imaging of cellular dynamics (61–64). Ongoing work continues to improve the technical capabilities of SRS such that real time imaging of fingerprint spectral features within live cells may be possible. Coherent anti-Stokes Raman scattering, or CARS - a similar contrast modality to SRS – has achieved considerably fast imaging throughput at high spectral resolution over the span of the CARS spectrum (5ms/px dwell times over >3000 cm^−1^ bandwidth) (61, 65). While this approach was too slow for spatially resolving cellular dynamics in real time for our study, broadband CARS approaches may be suitable for numerous other biological applications with different instrument performance needs.

Though the data presented here offer support for the involvement of lipid dynamics in INS, it needs to be noted that focus precompensation and hsSRS does not readily show the absolute magnitude of deformation in the cell membrane during INS. With a molecular dynamic model of INS biophysics, simple bilayer geometry simulations may enable some degree of calibration to correlate observed hsSRS spectra with lipid bilayer physical properties. Without clear approximations of lipid bilayer physical or electrical properties, it becomes difficult to judge or estimate the cell capacitance changes postulated to depolarize cells from SRS data alone. Integrating voltage imaging or electrophysiology alongside our existing hsSRS experimental preparation may be helpful in identifying a relationship between lipid dynamics and capacitance. Imaging systems with framerates exceeding 1 kHz can provide a window into these dynamics – however we were unable to reach such high framerates with our system without damaging cells. More generally, our results provide supportive evidence of the role lipids play in INS however, the data does not show a causal relationship between lipid dynamics and INS. Further, our imaging approach does not differentiate between extracellular membranes and intracellular organelle membranes. Transmembrane protein sensitivity to INS phenomena is still not clear, though it is known that different molecular pathways can be actuated depending on cell phenotype (20, 21, 66–70). Despite these caveats, the provided data clearly demonstrates that lipid bilayer dynamics are changing during INS and these changes track with magnitude of stimulus. These results provide validation of the current mechanism’s key assumptions in a live neural cell model. The understanding of this concept serves as a crucial basis for understanding of label free neuromodulation more broadly. Further, the general experimental framework presented here is readily applicable to other methods of directed energy neuromodulation as well as in the study of other dynamic processes.

The mechanistic basis of directed energy label-free neuromodulation has long been a question lacking complete answers (71, 72). Having a better understanding of how directed energy in the optical domain can be used to modulate brain function opens the door for innovation in neuromodulation to improve spatial targeting, temporal accuracy, and long-term utility, optically or otherwise. Extending these understandings to the development of new neuromodulation methods, neural prostheses, and therapeutic interventions provides a promising outlook for directed energy approaches. Whether the mechanistic basis for methods of directed energy neuromodulation, such as infrared, ultrasonic, or radio frequency-based approaches, are shared remains to be demonstrated. Our approach may serve as a valuable benchmark for answering such questions in the future as technology in neuromodulation and hsSRS imaging continues to develop.

## Conclusion

We have used hsSRS to experimentally demonstrate the mechanistic involvement of lipid dynamics in INS in live neural cells. Our results provide direct supportive evidence of lipid bilayer structural changes related to thermally induced *trans-gauche* isomerization of lipid tail hydrocarbon chains during INS. These experimental observations are in line with the currently proposed mechanistic model of INS. The implications from our results reinforce the idea that the photothermal basis of INS may be driving a general, nonspecific effect in live cells that evokes a multitude of physiological responses. The experimental framework also highlights the utility of hsSRS microscopy in addressing questions with high temporal resolution requirements and will continue to provide fruitful information about live cell biophysics beyond neuromodulation.

## Acknowledgements

The authors wish to thank Dr. Manqing Wang, Dr. Paul Stoddart, Dr. William Patrick Roach, Dr. Mark Hutchinson, and Dr. Valentina Benfenati for their discussion and guidance that formed the early basis for this work. The authors would also like to thank Dr. Bruce Tromberg for his suggestions and guidance for the ratiometric fluorescence imaging experiments presented in this study. The authors also thank Dr. Bryan Millis for his input on the manuscript.

## Author Contributions

AMJ, EDJ, and WRA conceived the idea for the manuscript. AMJ and EDJ secured funding support for the published work. AMJ, EDJ, GT, RG, and WRA designed the experiments. AL assisted in identifying, preparing, imaging the control samples for the study, and interpreting the results. AIBC assisted in preparing cell cultures, formulating experimental approaches, and data analysis. RG, AL, and GT contributed to data processing and analysis. BRJ, CD, AL, and WRA prepared the multilamellar vesicles. WRA assisted in all sample preparations, performed all imaging experiments, image processing, data analysis, data visualization, and wrote the manuscript. All authors contributed to editing manuscript.

## Data Availability

Any raw or processed data, processing, and analysis code are available upon request from the corresponding authors.

## Supplemental Figures

**Figure S1:**
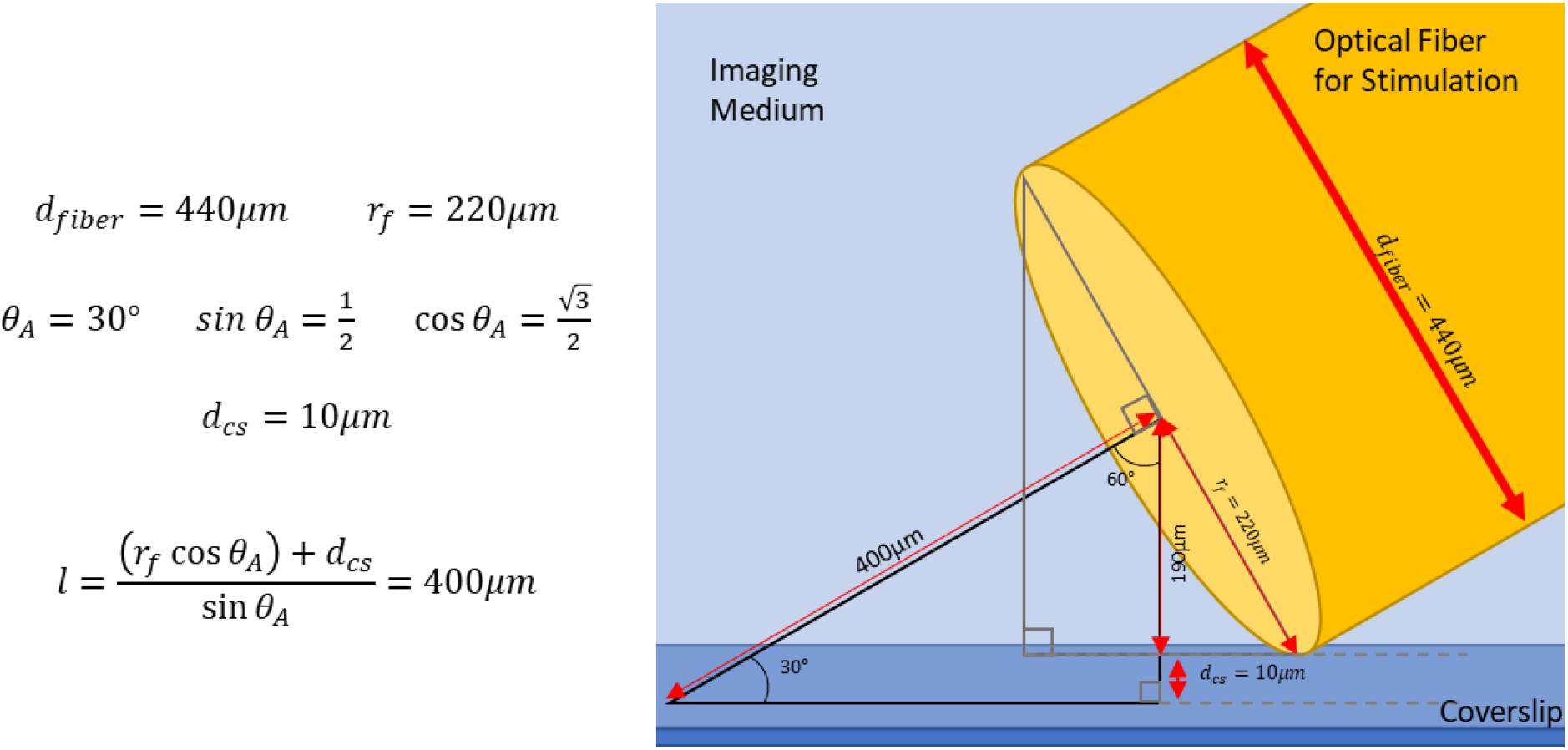
A) Illumination geometry and B) calculation of approximate fiber distance for estimating radiant exposure – where d_fiber_ is the optical fiber diameter, r_fiber_ is the optical fiber radius, θ_A_ is the fiber approach angle, d_cs+_ is the fiber edge’s distance off of the surface of the cover slip, and l is the normal distance from the optical fiber face to the cover slip plane.

**Figure S2:**
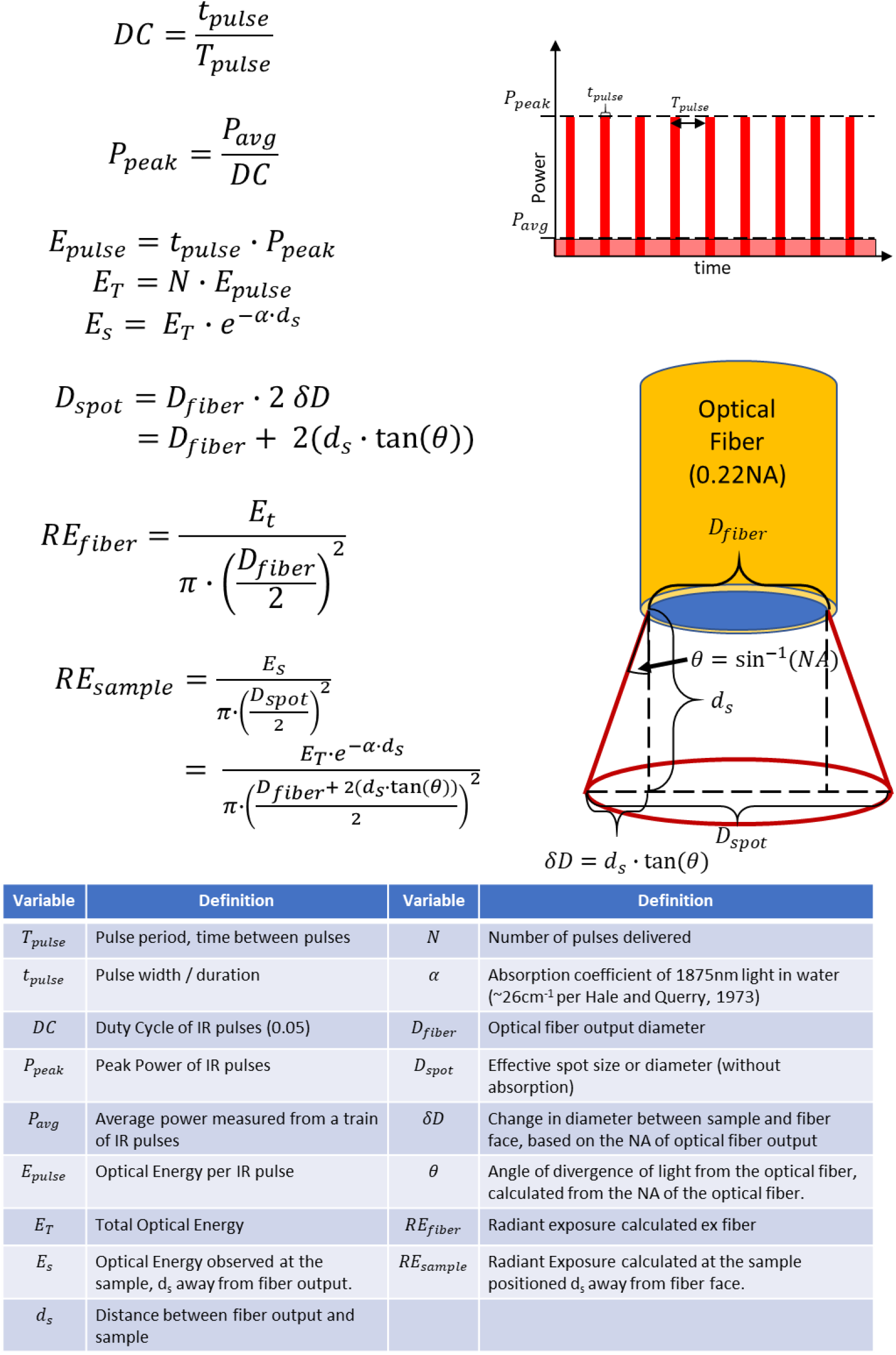
**Optical dosage calculations at the cell imaging plane based on an absorption-dominated photon distribution in homogenous medium, assuming negligible scattering and non-angled fiber approach to the sample**

**Figure S3:**
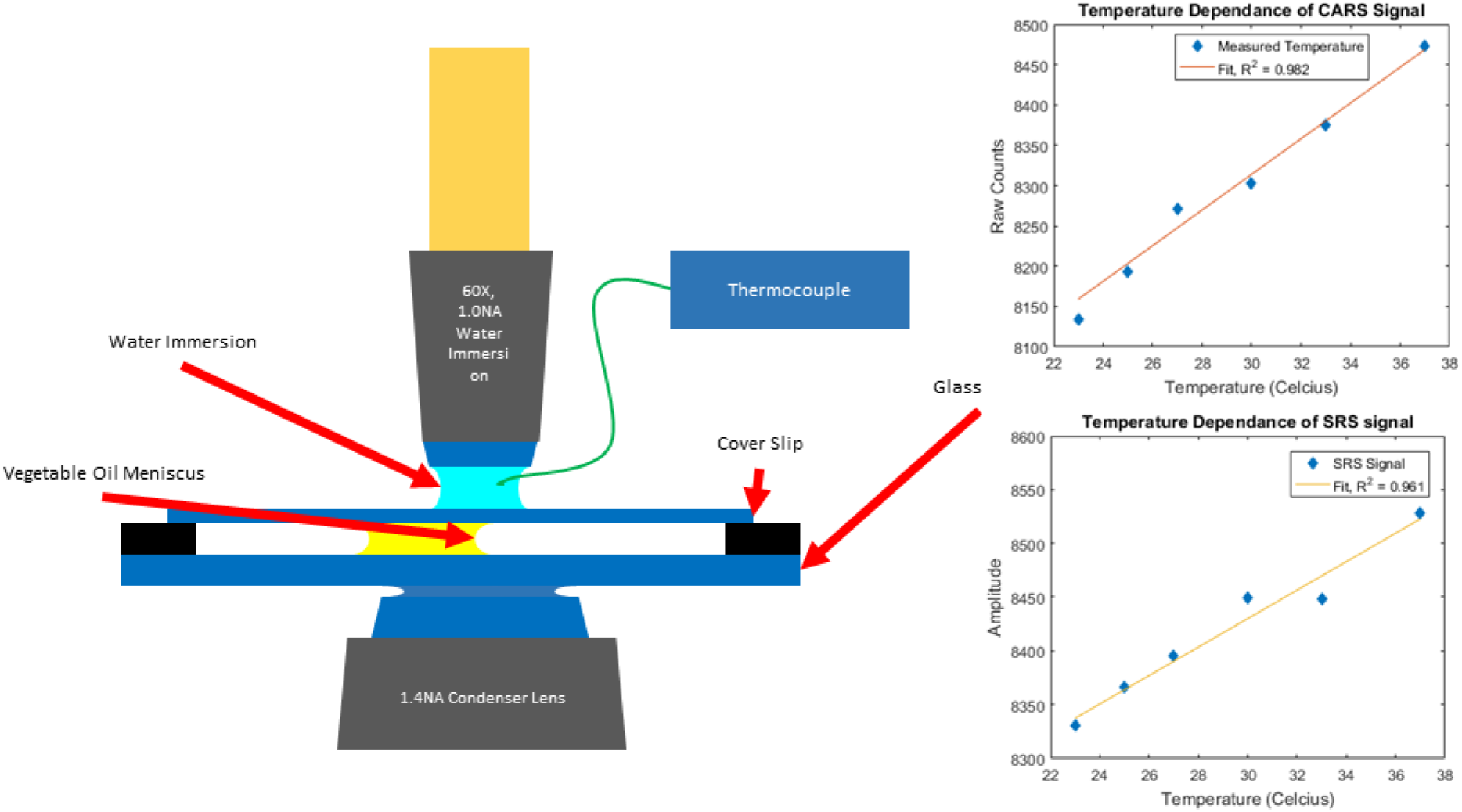
Temperature dependence of 2930 cm^−1^ CARS and SRS signal. A) experimental imaging and temperature measurement setup. B) Raw intensity measurements of vegetable oil meniscus as a function of temperature.

**Figure S4:**
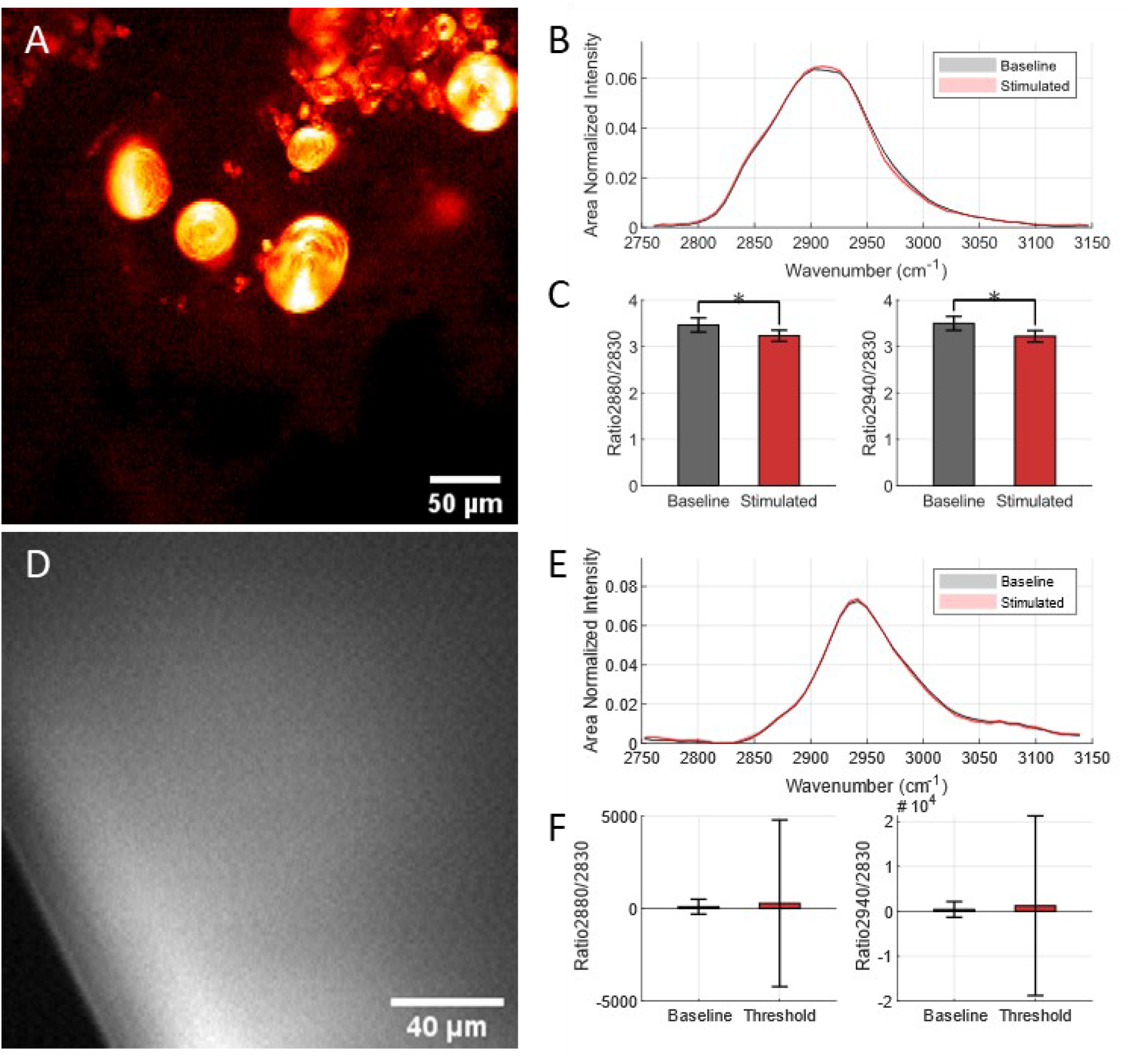
Validation of IR stimulated hsSRS images on isolated control sample preparations of major biological Raman scatterers. (A) SRS image of a 10% bovine serum albumin (BSA) sample in phosphate buffered saline as a control sample to measure protein SRS spectra (B) baseline and IR-stimulated SRS spectra observed in BSA solution. (D) SRS image of multilamellar vesicles at 2930 cm^−1^ resonance. (E) SRS spectra of baseline and IR-stimulated MLVs. (C, F) Ratiometric comparison of BSA and MLV SRS spectra, respectively, of resonances indicative of lipid membrane biophysical dynamics.

**Figure S5:**
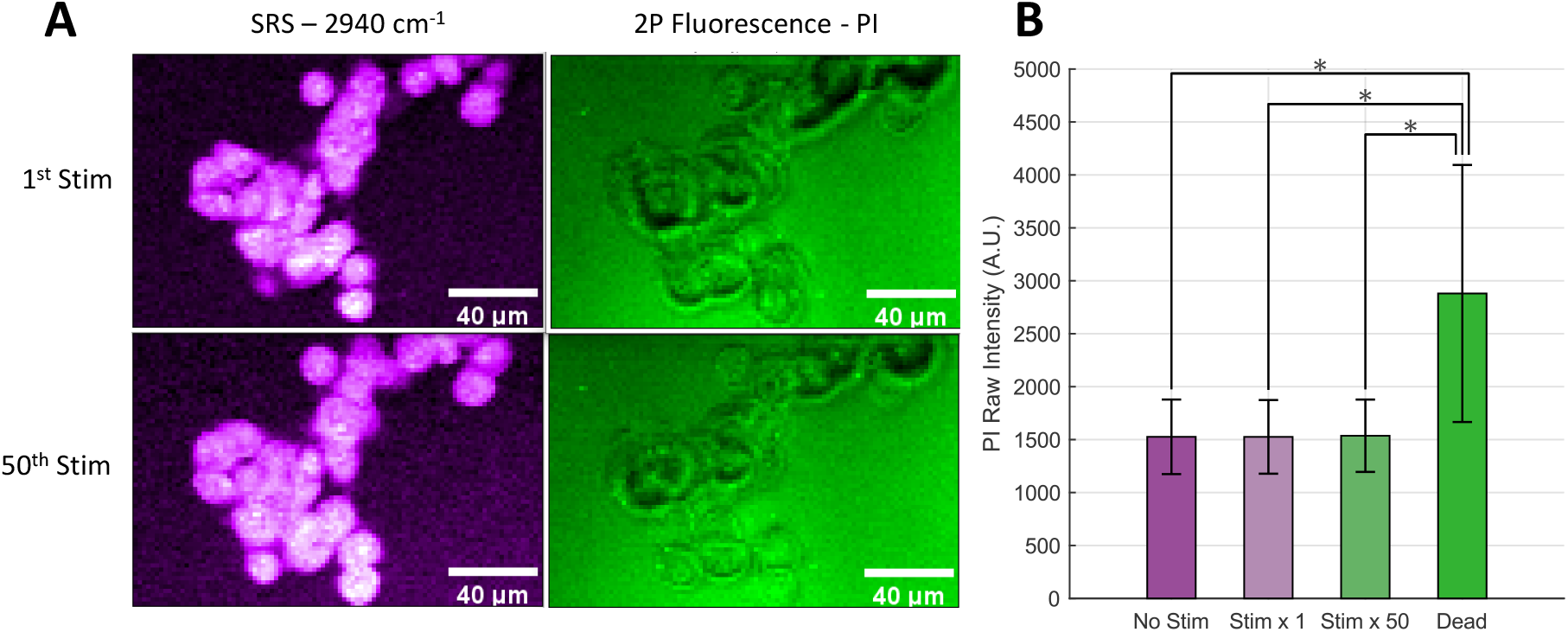
NG108 Cell Viability following hsSRS and repeated INS –. (A) Representative average intensity projection images of NG108 cells with SRS (left, magenta) and 2P fluorescence (green, right, identical intensity image scaling) of a cell viability indicator, propidium iodide (PI). Slight differences in cell morphology appear after 50 rounds of INS (bottom) compared to 1 round of INS (top). No substantial update of PI was observable. Scale bars are all 40 μm in width. (B) Intensity level comparison of PI fluorescence in cells exposed to different amounts of threshold INS events. No significant differences observed between non-stimulated and stimulated conditions. Significantly lower fluorescence compared to positive control of dead cells across all conditions. Asterisk indicates p < 0.05 based on a 2-sided student’s t-test comparisons of cell intensity means and standard deviations across all measured cells (n = 38).

**Figure S6:**
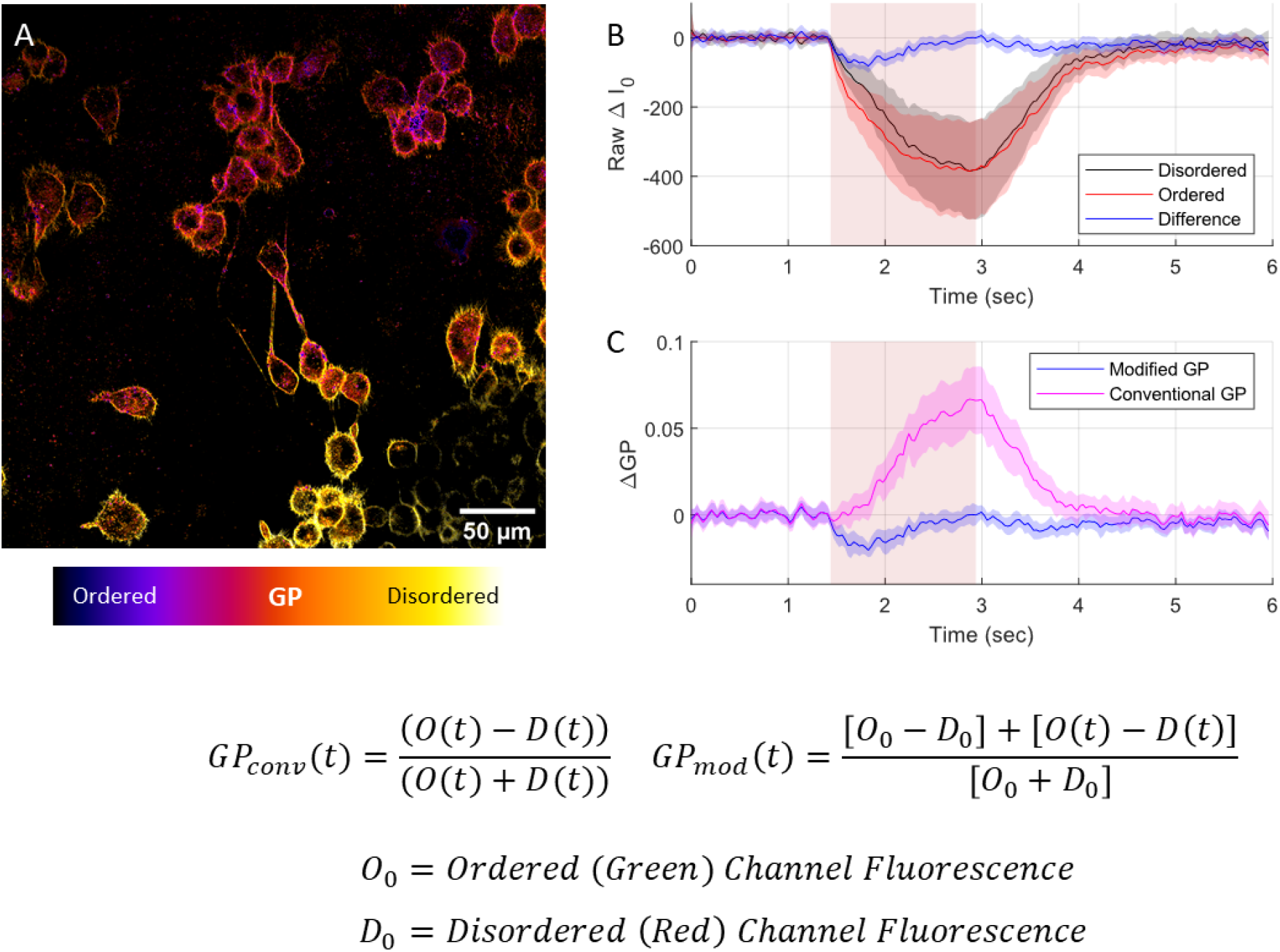
An intensity-invariant metric of general polarization for di-4-ANNEPS imaging of cells during IR stimulation, where signal loss from thermal lensing significantly impacts perceived signal interpretation. A) di-4-ANNEPS loaded NG108 cells. B) Baseline-offset mean detected intensities of mean disordered (black line) and ordered (red line) of all cells in a given experiment, plotted alongside the difference of detected intensities (Ordered – Disordered) C) Calculated conventional general polarization timeseries during IR stimulation alongside adapted general polarization calculation. D) Conventional and adapted GP metric calculations alongside each other. Eliminating the time dependance of the denominator term circumvents the defocusing artifact’s impact on the GP calculation.

